# Cross-compatibility of five highbush blueberry varieties and ideal crossing combinations

**DOI:** 10.1101/742114

**Authors:** Keanu Martin, Corneile Minnaar, Marinus de Jager, Bruce Anderson

**Affiliations:** Department of Botany and Zoology, Stellenbosch University, Matieland 7602, Cape Town, Western Cape, South Africa; Department of Conservation Ecology and Entomology, Stellenbosch University, Matieland 7602, Cape Town, Western Cape, South Africa

## Abstract

Blueberry plants require large quantities of pollen deposited on stigmas to produce commercial-quality fruit. Like many agricultural crops, the interaction between pollen-source variety and pollen-recipient variety can be a major determinant of fruit quality in blueberries. However, little information exists to guide growers in optimising fruit set and quality. Using five commonly grown blueberry varieties, I determined whether crossing between varieties (inter-varietal) increased fruit mass and decreased developmental time relative to crossing within a variety (intra-varietal), and if so, what the best crossing combinations are. While intra-varietal pollination often produced fruit, for certain varieties the fruit set and fruit mass were highly reduced compared to inter-varietal pollination. Furthermore, intra-varietal pollination resulted in longer fruit developmental time in comparison to pollination between varieties. For the same pollen-recipient variety, inter-varietal crosses typically outperformed intra-varietal crosses in fruit mass and developmental time; however, the extent to which inter-varietal crosses outperformed intra-varietal crosses differed between pollen-donor varieties. This result suggests that combinations of varieties are not trivial as some inter-varietal combinations may outperform others. Furthermore, some varieties appear to be more susceptible to the negative effects of intra-varietal crosses than others and that less susceptible varieties may be better suited to conditions where pollinator movement is poor. While our study can guide growers in determining optimal co-planting schemes for the varieties tested, for example in South Africa where these varieties are frequently grown. It also serves as a blueprint for similar compatibility studies that can easily be performed prior to planting to determine the best inter-varietal combinations.

## Introduction

Agriculture is continually focused on improving yields, but crops are persistently barraged with a range of problems including pests (Isman, 2000; Kumar et al., 2008), lack of nutrients (Parikh and James, 2012), and droughts (Bodner et al., 2015), that may reduce yield or even induce crop failure (Mendelsohn, 2007). Growers therefore persistently seek innovative ways to improve pest-control, plant nutrition, and irrigation to overcome these problems and increase yields (Bodner et al., 2015; Isman, 2000; Kumar et al., 2008; Mueller et al., 2012). Yet, even when fertigation and pest-control are ideally managed, many crops may still fail to produce optimal yields if they are not adequately pollinated (Broussard et al., 2017; Fijen et al., 2018). Growers of animal-pollinated crops therefore need to manage an additional dimension of crop production—pollination services—to ensure that enough of the correct pollen lands on recipient stigmas (Eaton and Nams, 2012; Gaines-Day and Gratton, 2016; Stern et al., 2001).

The simplest intervention to increase crop yields through pollination is to increase pollinator numbers (Eaton and Nams, 2012; Gaines-Day and Gratton, 2016; Stern et al., 2001). For example, Stern *et al*., (2001) found doubling honey bee hives caused an increase in the number of honey bee foragers leading to increased fruit set and yield of ‘Red Delicious’ apple trees by 50–100%. However, increasing hive numbers does not always result in greater yields: Eaton and Nams., (2012) found that increasing bee hive density caused an increase in fruit set and yield in blueberries but only to a point. This suggests that at a certain level of abundance, further investment to increase pollinator numbers may be met with diminishing returns on yield (Thomson, 2003) (Fig. 1.1). Thus, although increasing pollinator numbers is often one of the first means employed to increase low fruit yields, it is easy to over-shoot on this solution and it is important to understand where and when the point of diminishing returns is reached.

**Fig 1.1:**
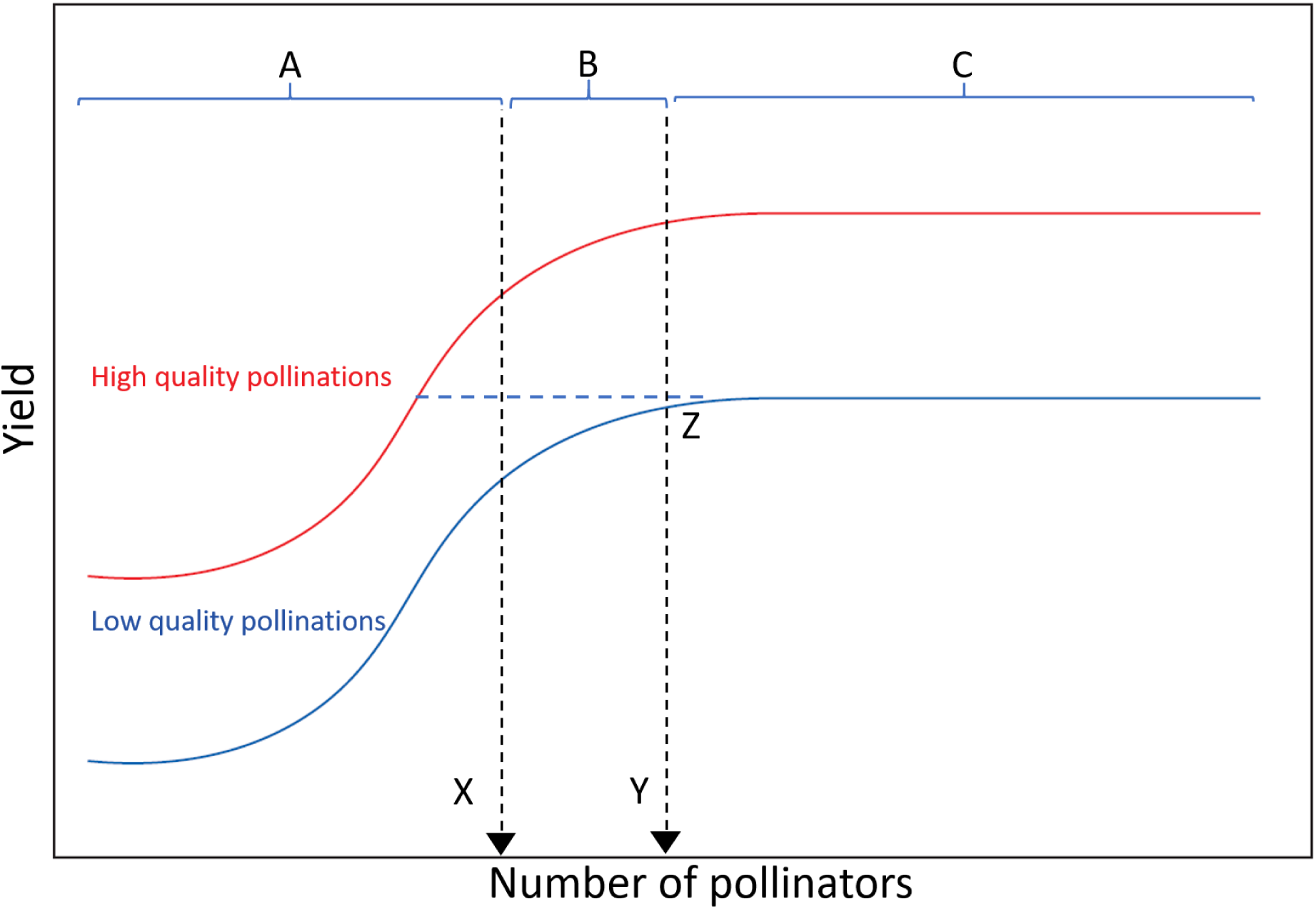
Hypothetical relationship between fruit yield and the quantity of pollinators for high and low pollination quality. A) Increasing pollinator numbers is likely to have a marked positive effect on yields until most stigmas are saturated with pollen (point X). Thereafter, additional increases in pollinator numbers are expected to progressively result in smaller and smaller yield increases until all stigmas are saturated with pollen (point Y). The area between points X and Y (B) is the area of diminishing returns with increased investment in pollinators. C) After this point, the addition of more pollinators will not increase yields, however yields can be increased by increasing the quality of pollen receipt (red curve versus blue curve). In fact, increases in pollen quality have the potential to increase yields across the entire spectrum of pollinator numbers. Furthermore, low numbers of pollinators, carrying high quality pollen may generate similar yields to high numbers of pollinators carrying low quality pollen (dotted line Z).

The second method of increasing crop yields through pollination is to improve the quality of pollen reaching stigmas, rather than the quantity. In many crops, fruit quality and yield vary depending on the source of pollen, i.e., pollen from the same variety (intra-variety pollination) or pollen from different varieties (inter-variety pollination). For example, ‘Red delicious’ apples require inter-varietal pollination to produce fruit (Stern et al., 2001) as most varieties are self-incompatible.

Similarly in dates, fruit set, weight, maturity and marketable yield are all affected by pollen source (Rezazadeh et al., 2013). Importantly, increasing the quality of pollen can have positive effects on yields, even when further increases in pollinator numbers have negligible effects on yield (Fig 1).

Blueberry fruit quality is directly affected by pollen source, where inter-varietal pollination often positively affects fruit size, weight, and developmental time when compared to pollination within varieties (Bell et al., 2012; Dogterom et al., 2000; Ehlenfeldt, 2001; Gupton and Spiers, 1994; Harrison et al., 1993; Huang et al., 1997; Müller et al., 2013; Taber and Olmstead, 2016; Usui et al., 2005). In response to this constraint, growers often plant different varieties in adjacent rows in an orchard in attempt to increase yield through inter-varietal pollination. While inter-varietal pollination is likely to increase yield, varieties may differ in their compatibility with each other. Knowing which varieties should be planted together is thus of crucial importance to growers. Despite this, very few multi-variety compatibility studies exist to guide growers in deciding which varieties to co-plant.

In this paper, I investigate the interactive effects of pollen origin and pollen recipient on fruit yields for some of the most commonly planted blueberry varieties, especially in South Africa. This will help blueberry growers to optimize inter-varietal planting beyond simply planting two different varieties together. First, I determined whether inter-varietal pollination generally increases fruit mass relative to intra-varietal pollination. I then asked whether there were ideal combinations of varieties which maximize fruit production and yield. I hypothesized that inter-varietal pollination would result in significantly larger yields than crossing within varieties. I also expect that yields will be significantly affected by the choice of recipient and donor pairs.

## Methods and materials

### Study site

This study was conducted in an experimental blueberry plot in Klapmuts (Western Cape, South Africa, 33°49′42.5″S 18°55′03.5″E). The experimental plot was stocked at a density of 10 honey bee hives per hectare.

### Varieties and experimental design

I conducted a series of hand pollinations between concurrently flowering blueberry varieties, Emerald, Eureka, Snowchaser, Suziblue and Twilight. Emerald and Eureka were the first varieties to bloom, followed by Snowchaser, Suziblue and the Twilight. Pollination for each variety was conducted at 25% bloom. Therefore, varieties such as Emerald and Eureka could be pollen donors to all varieties but not recipients to all varieties. Each possible pollination combination for each variety (i.e., crosses within varieties and between varieties that overlap in flowering period, see Table 1.1) was replicated across nine plants, with each pollination combination repeated three times per individual plant. This yielded a total of 27 replicates per pollination combination. Since blueberry plants of the same variety are clones, I did not expect variation between plants to strongly affect our results. In addition, by using fewer individual plants, but with repetition within plants, I was able to conduct my experiments across varieties within a small unit of space (experimental plot size: 50 m X 115 m). This approach reduced variation in external factors that may influence fruit development (e.g. soil nutrition, moisture and irrigation, and exposure to sunlight and wind) that typically vary in situ across large-scale multi-variety plantings.

**Table 1.1:**
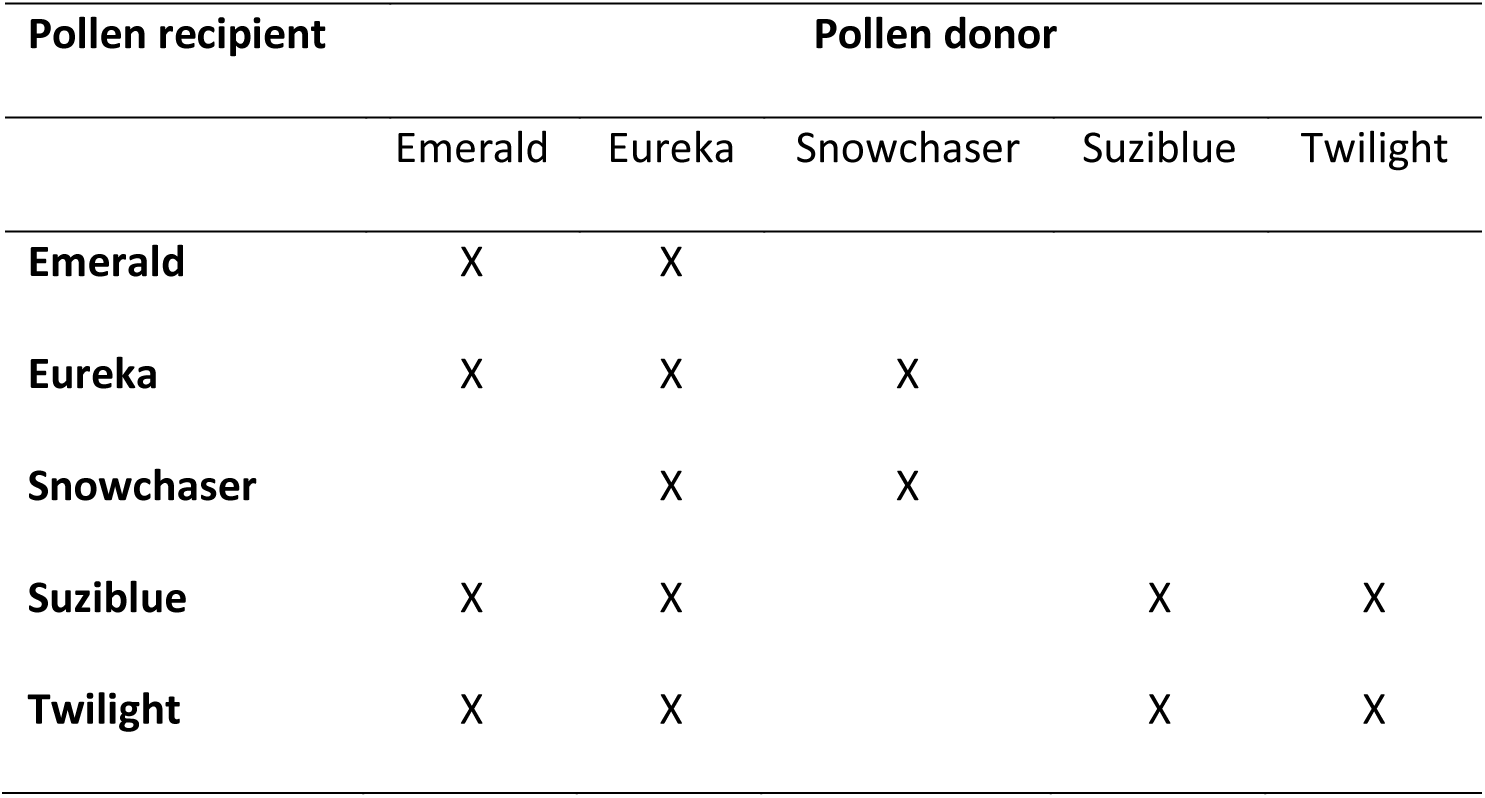
Cross-combinations of pollination donor–recipient pairs used in this study. Blank cells indicate varieties that do not overlap in flowering time. Emerald and Eureka were the first varieties to bloom, followed by Snowchaser, Suziblue and the Twilight. All pollinations were conducted when a variety was at 25% flowering. Varieties could therefore be donors to later flowering varieties but not recipients.

### Hand-pollination protocol

To ensure that experimental flowers were virgins, buds that were about to open were bagged three days before hand pollination. The day before pollination, I collected pollen from pollen donor flowers (approximately five flowers per pollen application) (Dogterom et al., 2000). Pollen was extracted from donor flowers by removing the corolla with a scalpel and agitating the anthers with forceps, causing the poricidal anthers to release pollen that I collected in a Petri dish. Prior to hand-pollination, recipient flowers were marked and emasculated. I did this by removing all stamens with a pair of fine forceps, ensuring that no self-pollination could take place. After removing anthers, I checked stigmas with a 20x hand lens to ensure no pollen deposition occurred. I mixed pollen from donor flowers on different individual plants to ensure that recipient flowers received pollen from multiple donors. I dipped recipient stigmas into this pollen mix and visually confirmed that the stigma was saturated with pollen. To prevent honey bees from depositing additional pollen of unknown origin onto the stigma I placed a fine mesh bag over hand-pollinated flowers following hand-pollination. Once flowers wilted, I removed their bags.

### Fruit quality variables

After hand pollinations were conducted, I checked fruit development weekly to determine whether fruit were mature (the entire fruit had turned a uniform dark blue) and ready for fruit-quality measurements. I harvested and weighed mature fruit by hand. I also measured the diameter of each fruit on the day it was harvested. The developmental time for each fruit (the number of weeks from pollination to harvesting) for each donor–recipient pair was also determined. Similar to fruit mass and diameter, the developmental time is a significant determinant of fruit value, because early fruit are more valuable than late fruit, since early fruit can be sold at higher prices when market demand is not saturated (Dogterom et al., 2000; Sobekova et al., 2013). I also calculated the percentage fruit set per donor–recipient pair. I combined fruit mass and percentage fruit set for donor-recipient pair *i* to calculate the adjusted fruit mass (i.e. a proxy for realised yield) for donor–recipient pair *i* as follows:

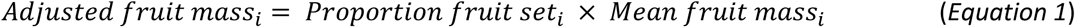

### Statistical analyses

I used linear mixed-effects models with fruit mass, diameter, developmental time and adjusted fruit mass as a function of donor–recipient pair (fixed effect). I allowed the model to vary in both slope and intercept (without covariance) per plant ID to account for variation among individual plants (plant ID as a random effect), thus allowing the model more sensitivity to detect differences in fruit quality measures as a result of donor–recipient pair in isolation. Linear contrasts (Tukey) were used to test differences between donor–recipient pairs. To assess model fit to our data, I used Nakagawa *R*^*2*^ values (Nakagawa and Schielzeth, 2013), which provides both conditional variance (*R*^*2*^*c*) and marginal variance (*R*^*2*^*m*) estimates for linear mixed-effects models that can be equated to traditional *R*^*2*^values. Conditional *R*^*2*^ values (*R*^*2*^*c*) reflect the variance explained by the entire model (fixed effects and random effects), whereas marginal *R*^*2*^ values (*R*^*2*^*m*) indicate the variance explained by the fixed effects alone.

### Inter-vs. intra-varietal pollination

To determine whether inter-varietal pollination was beneficial to blueberry production, I grouped inter-varietal pollination combinations by recipient variety and compared overall fruit quality resulting from inter-varietal pollination to intra-varietal pollinations for that variety. I used the same linear mixed-effects model approach for this comparison, but with adjusted fruit mass as a function of intra-vs inter-varietal pollination (fixed effect). As above, I allowed the model to vary in both slope and intercept (without covariance) per plant ID to account for variation among individual plants (plant ID as a random effect). Since there were only two levels for the fixed effect, I compared model estimates for intra-vs inter-varietal pollination directly and assessed model fit to our data using Nakagawa *R*^*2*^ values (Nakagawa and Schielzeth, 2013).

All statistical analyses were conducted in R (version 3.3.2) (R Core Team, 2017) using the packages nlme (Pinheiro et al., 2017), multcomp (Hothorn et al., 2008), ggplot2 (Wickham, 2009), sjPlot (Lüdecke, 2017), car (Fox and Weisberg, 2011), lme4 (Bates et al., 2015), grid (R Core Team, 2017), gridExtra (Auguie, 2017), lattice (Deepayan, 2008), MuMIn (Barton, 2017), plyr (Wickham, 2011) and plotrix (Lemon, 2006).

## Results

### Inter-vs. intra-varietal pollination

Inter-varietal pollination resulted in larger fruit for Suziblue and Twilight, but not for Emerald, Eureka and Snowchaser. Inter-varietal pollination consistently resulted in a heavier adjusted fruit mass compared to intra-varietal pollination. Developmental time was largely unaffected by pollen source, except in the case of Suziblue and Twilight. Below, I discuss the results of our experiments in depth.

Across all varieties, inter-varietal pollination significantly (z=4.3580, p<0.001; Fig 1.2) increased adjusted fruit mass by 59% from 1.4943 ± 0.0828 g (mean ± SE) for intra-varietal pollinations to 2.3716g ± 0.1794 (mean ± SE) for inter-varietal pollinations.

**Fig 1.2:**
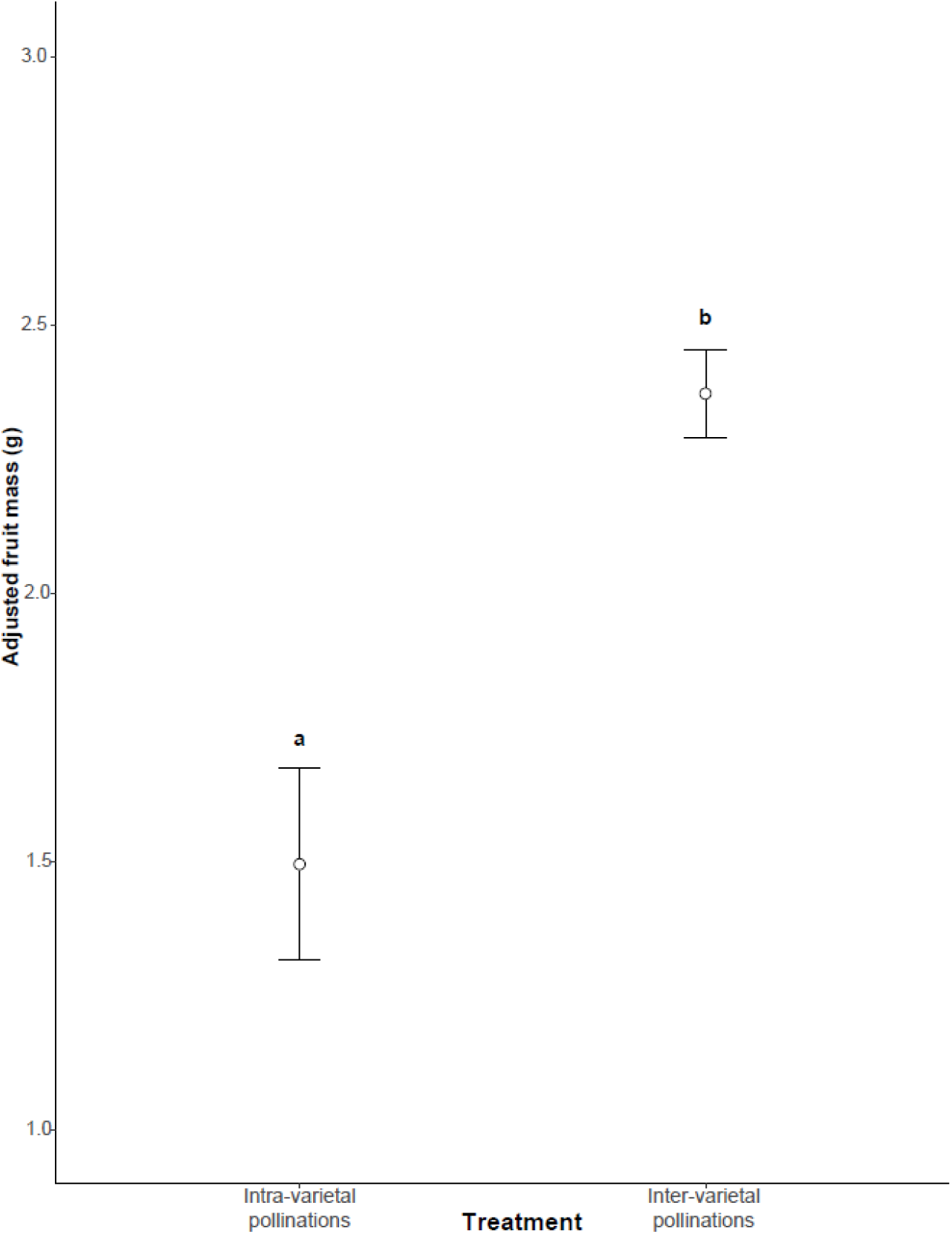
The overall effect of intra-varietal pollinations versus inter-varietal pollinations on adjusted fruit mass for five blueberry varieties. Adjusted fruit mass is the product of proportion fruit set and mean fruit set. Letters indicate significance (p<0.05) of linear contrasts (Tukey HSD). Error bars indicate standard error.

### Fruit quality from all pollination combinations per variety

Donor–recipient pairs explained most of the variance in developmental time compared to plant ID, since the conditional variance for the developmental time model was *R*^*2*^*c*=0.2905 (fixed- and random-effects). This was very similar to the marginal variance (fixed-effects only: *R*^*2*^*m*=0.2755). Emerald pollen shortened developmental time to 11 weeks ± 0.62 weeks (mean ± SE weeks) substantially (*z*= −3.8972, *p*<0.001; see Table S1) compared to intra variety pollen (Suziblue) (mean ± SE weeks = 13 weeks ± 0.4 weeks; Fig 1.3). Pollinating with Twilight pollen (*z*= 0.3880, *p*=0.9801; see Table S1) and Eureka pollen (*z*= −0.2760, *p*=0.9926; see Table S1) resulted in the same developmental time as within variety pollination (mean ± SE weeks = 13 weeks ± 0.36 weeks, mean ± SE weeks = 13 weeks ± 0.55 weeks respectively; Fig 1.3).

**Fig 1.3:**
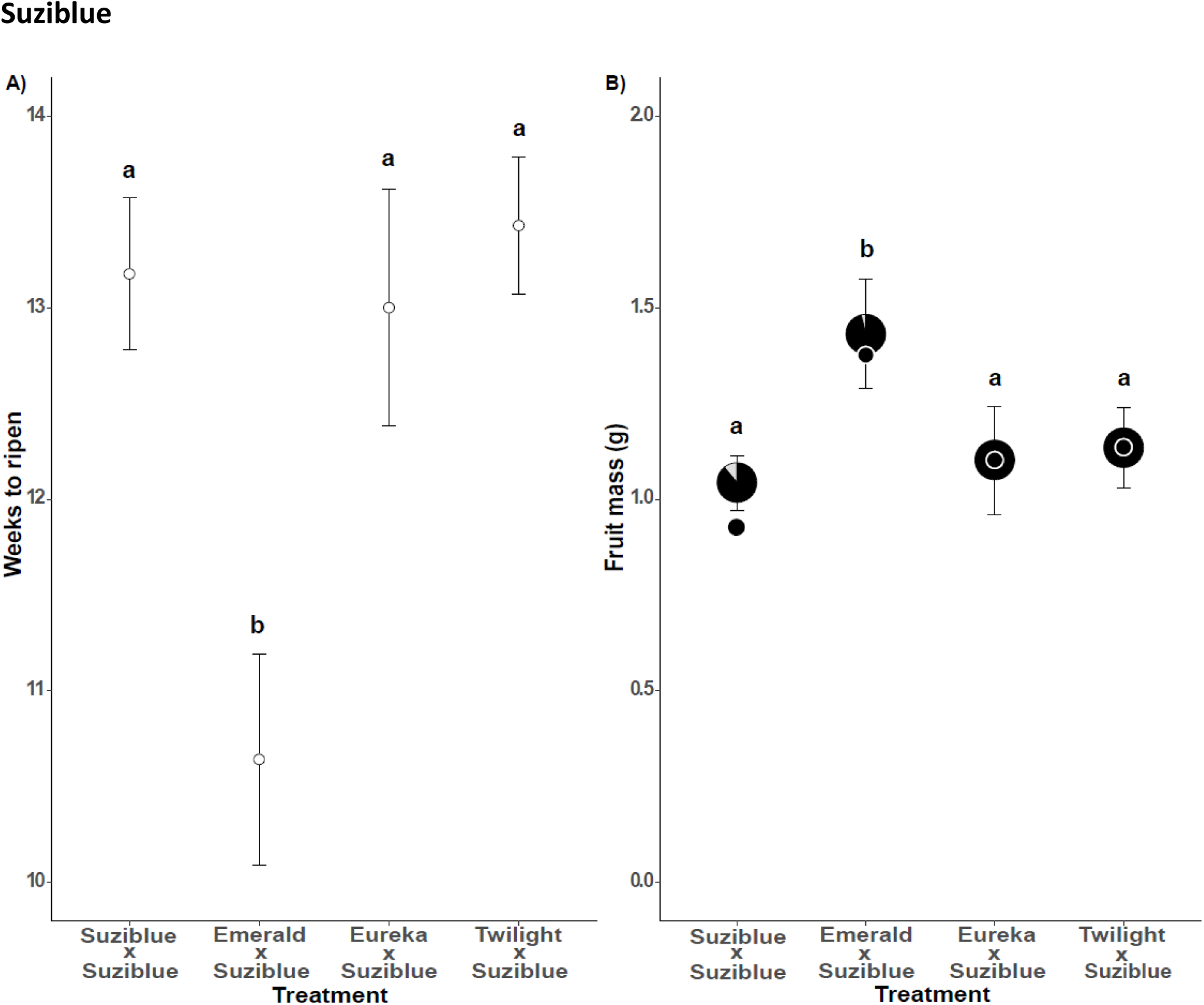
A) Mean number of weeks needed for the fruit to ripen for each treatment. B) Two measurements of fruit mass. Mean ± SD mass per fruit (pie charts ± whiskers) and adjusted fruit mass (Black dots). Black parts of pie charts also indicate the proportion fruit set for each treatment. Adjusted fruit mass is the product of proportion fruit set and mean fruit set. Letters indicate significance (p<0.05) of linear contrasts (Tukey HSD). Error bars indicate standard error.

Plant ID explained most of the variance in fruit mass compared to donor–recipient pairs, since the conditional variance for the fruit mass model was *R*^*2*^*c*=0.6942 (fixed- and random-effects). This was largely different to the marginal variance (fixed-effects only: *R*^*2*^*m*=0.1927). Inter-varietal pollinations using Emerald pollen significantly increased (*z*= 3.8720, *p*<0.001; see Table S1) the fruit mass of Suziblue by 38% to (mean ± SE fruit mass = 1.43 g ± 0.14 g). I did not detect a significant difference in fruit mass for Eureka x Suziblue (*z*= 0.5940, *p*=0.9322; see Table S1) and Twilight x Suziblue (*z*= 1.9910, *p*=0.1862; see Table S1) compared to intra-varietal pollination, with the average (± SE) fruit mass resulting from inter-varietal pollinations with Eureka (1.10 g ± 0.14 g) and Twilight (1.13 g ± 0.10 g) only 6% and 9% higher respectively than for Suziblue x Suziblue pollinations (1.04 g ± 0.07 g; Fig 1.3).

Donor–recipient pairs explained most of the variance in developmental time compared to plant ID, since the conditional variance for the developmental time model was *R*^*2*^*c*=0.4898 (fixed- and random-effects), and this was different to the marginal variance (fixed-effects only: *R*^*2*^*m*=0.3736). Emerald pollen and Snowchaser pollen resulted in a significantly shorter developmental time of 10 weeks ± 0.2 weeks (mean ± SE weeks; *z*= −6.0650, *p*<0.001; see Table S2) and 10 weeks ± 0.22 weeks (mean ± SE weeks; *z*= −5.5930, *p*<0.001; see Table S2), respectively. This was on average 2 weeks shorter than pollinating Twilight with its own pollen which had a ripening time of 12 weeks ± 0.43 weeks (mean ± SE weeks). I was unable to detect any difference in developmental time between pollinating Twilight with other Twilight pollen or Eureka pollen as Eureka also had a developmental time of 12 weeks (*z*= −0.6950, *p*=0.8990; see Table S2).

Plant ID explained most of the variance in fruit mass compared to donor–recipient pairs, since the conditional variance for the fruit mass model was *R*^*2*^*c*=0.3132 (fixed- and random-effects). This was largely different to the marginal variance (fixed-effects only: *R*^*2*^*m*=0.1375). Pollinating Twilight plants with pollen from other varieties significantly increased fruit mass in all donor–recipient pairs except Eureka (*z*= 1.503, *p*=0.4299; see Table S2; Fig 1.4). Pollinating Twilight with Emerald pollen resulted in the greatest fruit mass at 3.1 g ± 0.46 g (mean ± SE fruit mass) compared to Twilight plants pollinated with other Twilight pollen, a ca. 100% increase (*z*=2.7940, *p*=0.0259; see Table S2). Pollinating Twilight with Suziblue pollen also substantially increased fruit mass by 70% (*z*= 2.8770, *p*=0.0202; see Table S2) to 2.7 g ± 0.20 g (mean ± SE fruit mass).

**Fig 1.4:**
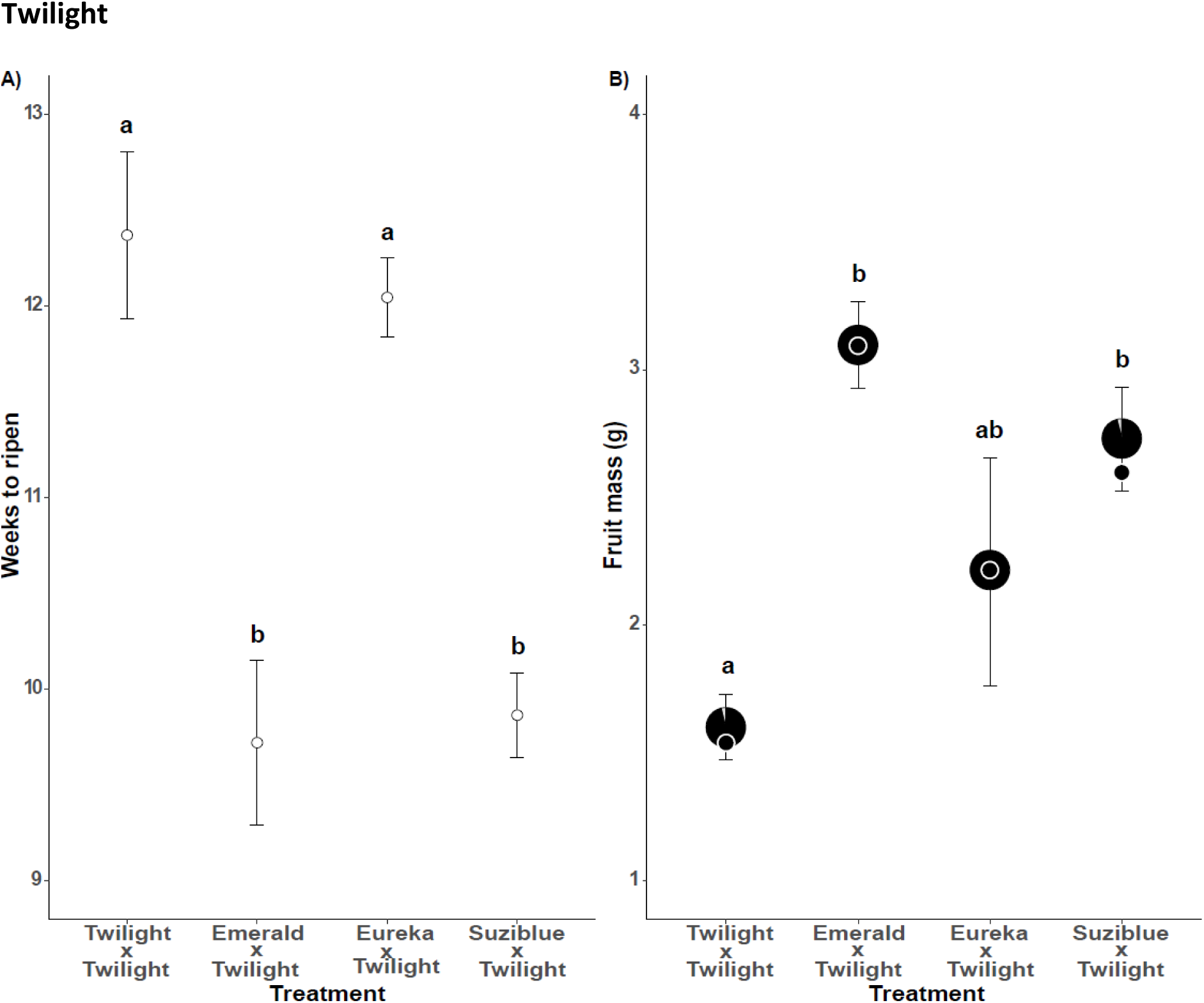
A) Mean number of weeks needed for the fruit to ripen for each treatment. B) Two measurements of fruit mass. Mean ± SD mass per fruit (pie charts ± whiskers) and adjusted fruit mass (Black dots). Black parts of pie charts also indicate the proportion fruit set for each treatment. Adjusted fruit mass is the product of proportion fruit set and mean fruit set. Letters indicate significance (p<0.05) of linear contrasts (Tukey HSD). Error bars indicate standard error.

Plant ID explained most of the variance in developmental time compared to donor–recipient pairs, since the conditional variance for the developmental time model was *R*^*2*^*c*=0.3132 (fixed- and random-effects). This was largely different to the marginal variance (fixed-effects only: *R*^*2*^*m*=0.0027). The lack of influence of donor–recipient pair (fixed effect) on developmental time is further supported by linear contrasts, which revealed no significant difference between donor–recipient pairs (*z*= −0.3570, *p*=0.7210; see Table S3). However, the average (± SE) developmental time resulting from inter-varietal pollination with Eureka (12 weeks ± 0.76 weeks) was one week shorter than Emerald x Emerald pollinations (13 weeks ± 0.59 weeks; Fig 1.5)—the direction (shorter developmental times for inter-varietal pollination) is consistent with expectations.

Donor–recipient pairs explained all of the variance in fruit mass compared to plant ID, since the conditional variance for the fruit mass model was *R*^*2*^*c*=0.0179 (fixed- and random-effects), and the marginal variance (fixed-effects only: *R*^*2*^*m*=0.0179); however, donor-recipient pair still explained a very small proportion of the overall variation in fruit mass. While I was unable to detect a significant difference in fruit mass between these donor–recipient pairs (*z*=0.7761, *p*=0.4380; see Table S3), the average (± SE) fruit mass resulting from inter-varietal pollinations with Eureka (1.62 g ± 0.16 g) was 12% higher than for Emerald x Emerald pollinations (1.46g ± 0.15 g, Fig. 1.5). The direction of this difference (increased mass for inter-varietal pollination) is consistent with expectations.

**Fig 1.5:**
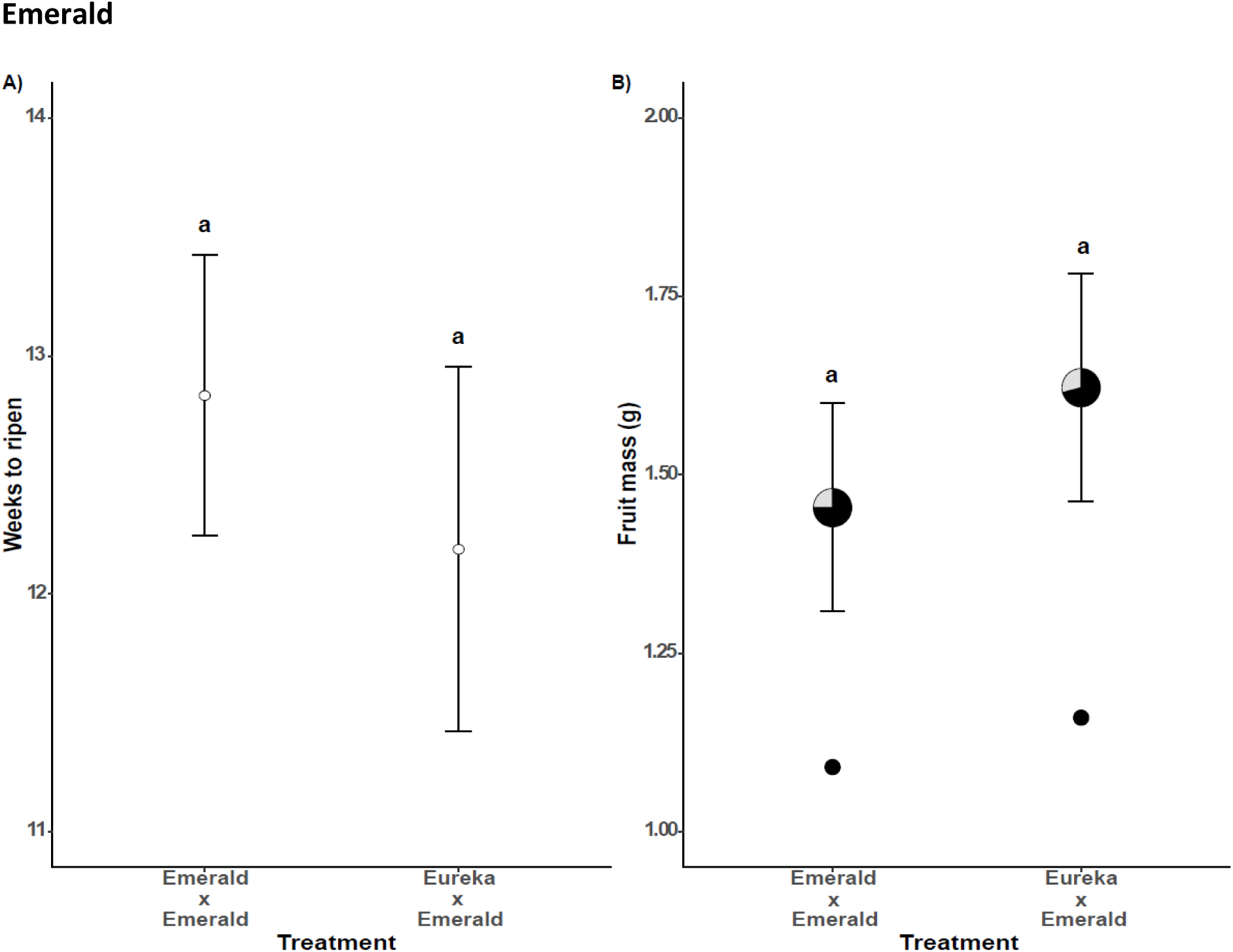
A) Mean number of weeks needed for the fruit to ripen for each treatment. B) Two measurements of fruit mass. Mean ± SD mass per fruit (pie charts ± whiskers) and adjusted fruit mass (Black dots). Black parts of pie charts also indicate the proportion fruit set for each treatment. Adjusted fruit mass is the product of proportion fruit set and mean fruit set. Letters indicate significance (*p*<0.05) of linear contrasts (Tukey HSD). Error bars indicate standard error.

Plant ID explained most of the variance in developmental time compared to donor–recipient pairs, since the conditional variance for the developmental time model was *R*^*2*^*c*=0.0209 (fixed- and random-effects). This was largely different to the marginal variance (fixed-effects only: *R*^*2*^*m*=0.0071). I was unable to distinguish a significant difference in developmental time between Emerald x Eureka (*z*= −0.3120, *p*=0.9480; see Table S4 and Snowchaser x Eureka (*z*=−0.6190, *p*= 0.8090; see Table S4). The average (± SE) developmental time resulting from inter-varietal pollinations with Snowchaser was 8 weeks ± 0.23 weeks and Eureka x Eureka pollinations resulted in a developmental time of 9 weeks ± 0.36 weeks (mean ± SE weeks; Fig 1.6). Pollinating Eureka with Emerald pollen also resulted in a developmental time of 9 weeks ± 0.26 weeks (mean ± SE weeks; Fig 1.6).

**Fig 1.6:**
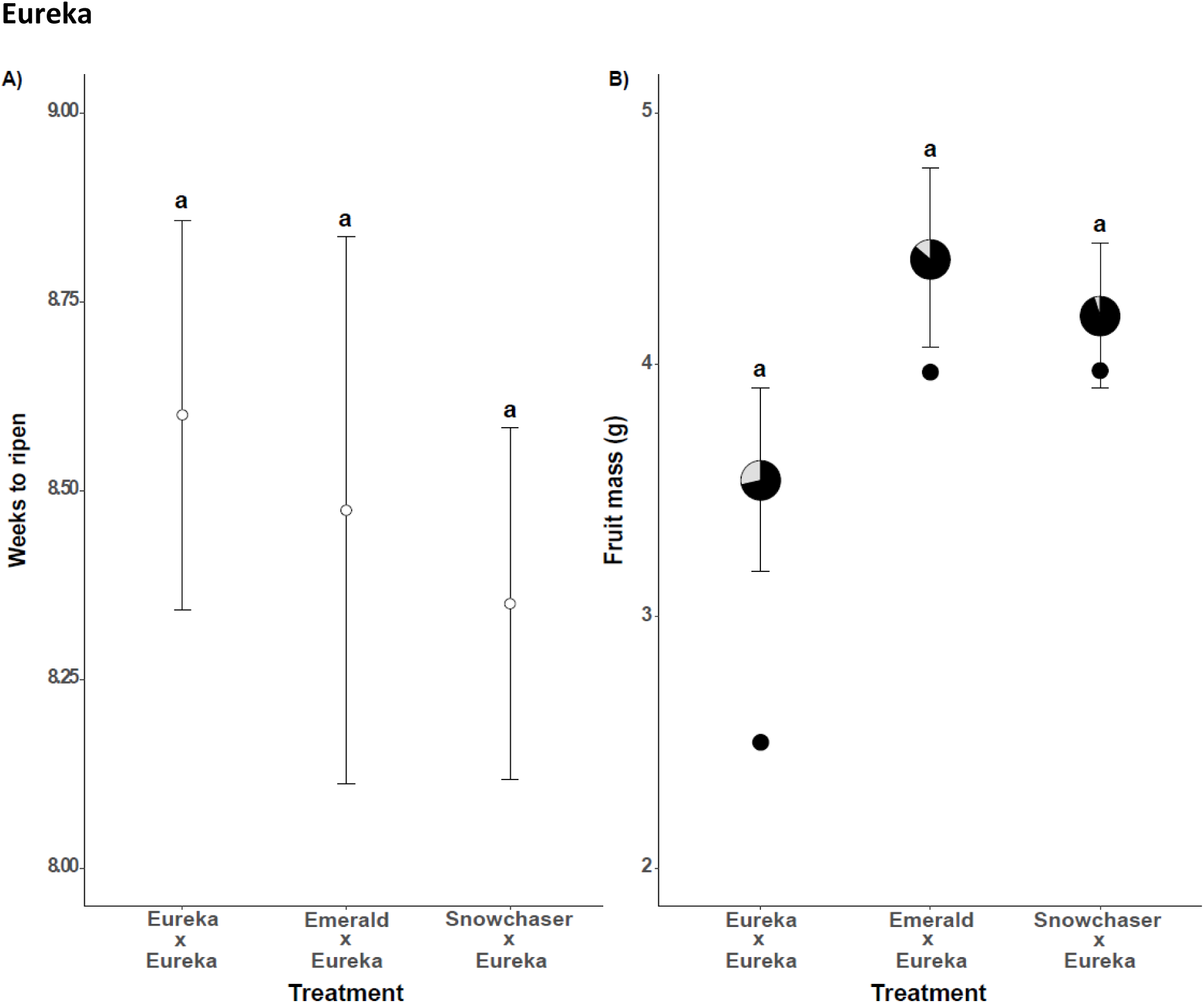
A) Mean number of weeks needed for the fruit to ripen for each treatment. B) Two measurements of fruit mass. Mean ± SD mass per fruit (pie charts ± whiskers) and adjusted fruit mass (Black dots). Black parts of pie charts also indicate the proportion fruit set for each treatment. Adjusted fruit mass is the product of proportion fruit set and mean fruit set. Letters indicate significance (p<0.05) of linear contrasts (Tukey HSD). Error bars indicate standard error.

Plant ID explained most of the variance in fruit mass compared to donor–recipient pairs, since the conditional variance for the fruit mass model was *R*^*2*^*c*=0.5211 (fixed- and random-effects). This was largely different to the marginal variance (fixed-effects only: *R*^*2*^*m*=0.0683). I was unable to detect a substantial difference in fruit mass between Emerald x Eureka (*z*=2.0960, *p*=0.0894; see Table S4) and Snowchaser x Eureka (*z*= 1.835, *p*= 0.1563; see Table S2). However, the average (± SE) fruit mass resulting from inter-varietal pollinations with Emerald (4.42 g ± 0.36g) and Snowchaser (4.19g ± 0.29g) was 25% and 18% higher respectively than for intra-varietal pollinations of Eureka (3.54 g ± 0.36g; Fig 1.6). Our level of replication may be the reason why the difference is too small to be statistically detected, however, the direction (increased mass for inter-varietal pollination) is consistent with our expectations.

Donor–recipient pairs explained all of the variance in developmental time compared to plant ID, since the conditional variance for the developmental time model was *R*^*2*^*c*=0.0179 (fixed- and random-effects), and the marginal variance (fixed-effects only: *R*^*2*^*m*=0.0179); however, donor-recipient pair still explained a very small proportion of the overall variation in developmental time. I could not detect a significant difference in developmental time between these donor–recipient pairs (*z*=−1.0860, *p*= 0.2780; see Table S5), the average (± SE) developmental time resulting from inter-varietal pollinations with Eureka was 10 weeks ± 0.16 weeks and for Snowchaser x Snowchaser pollinations the average (± SE) developmental time was 11 weeks ± 0.17 weeks (Fig 1.7).

**Fig 1.7:**
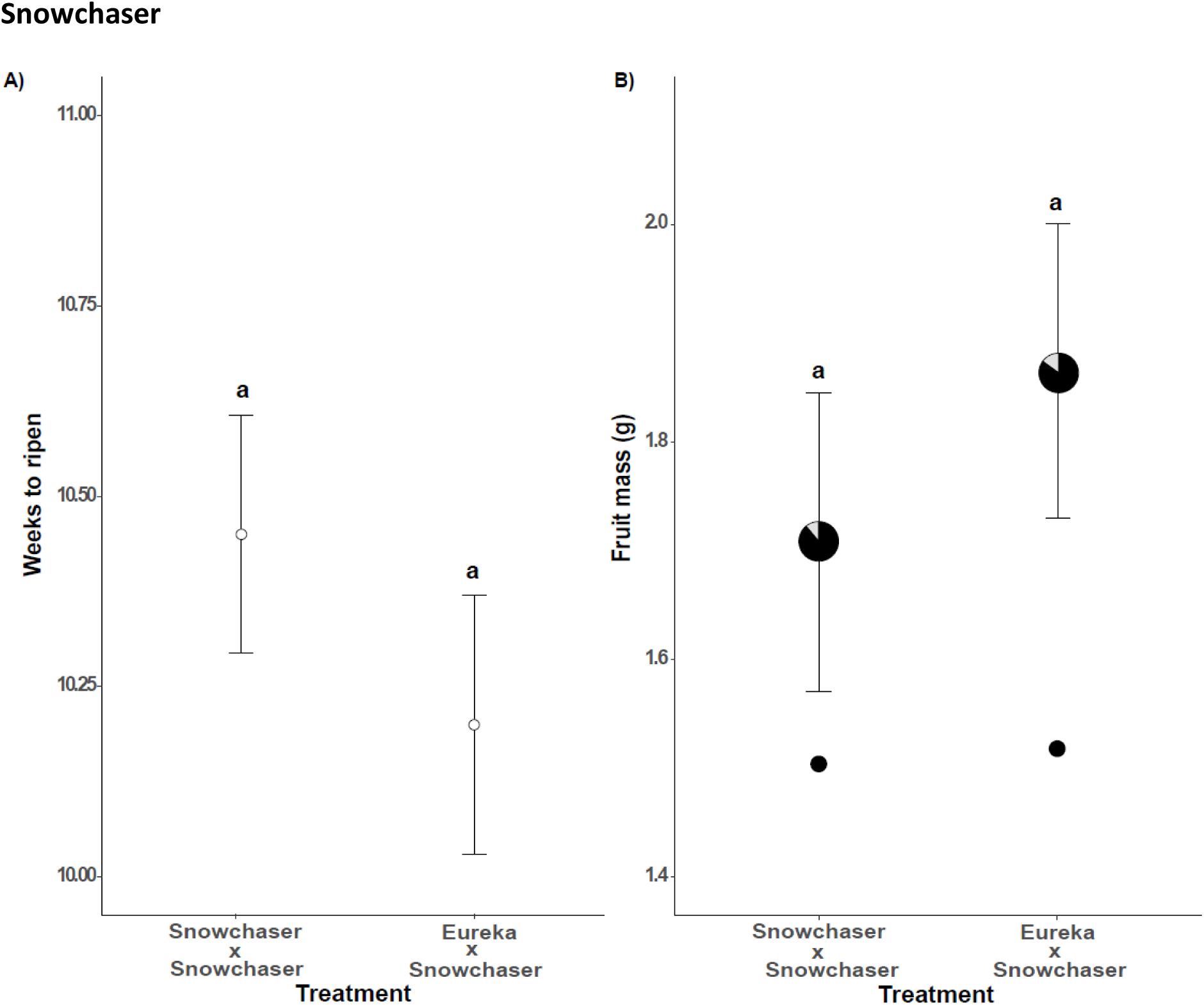
A) Mean number of weeks needed for the fruit to ripen for each treatment. B) Two measurements of fruit mass. Mean ± SD mass per fruit (pie charts ± whiskers) and adjusted fruit mass (Black dots). Black parts of pie charts also indicate the proportion fruit set for each treatment. Adjusted fruit mass is the product of proportion fruit set and mean fruit set. Letters indicate significance (p<0.05) of linear contrasts (Tukey HSD). Error bars indicate standard error.

Plant ID explained most of the variance in fruit mass compared to donor–recipient pairs, since the conditional variance for the fruit mass model was *R*^*2*^*c*=0.2376 (fixed- and random-effects). This was largely different to the marginal variance (fixed-effects only: *R*^*2*^*m*=0.0188). The average (± SE) fruit mass resulting from inter-varietal pollinations with Eureka (1.87 g ± 0.14 g) was 9% higher than for Snowchaser x Snowchaser pollinations (1.71 g ± 0.14 g), as expected reduced mass for intra-varietal pollination); however, my statistical analyses showed no significant difference between these donor–recipient pairs (z=0.9760, *p*=0.3290; see Table S5).

## Discussion

This study revealed a general trend of crosses within varieties producing lower adjusted fruit masses than crosses between varieties. Furthermore, differences in the source of pollen can affect both fruit mass and developmental time of multiple blueberry varieties. Inter-varietal pollination significantly increased fruit mass for Suziblue and Twilight and decreased developmental time. These differences in fruit mass among inter-varietal pollinations have real-world implications for growers and underscores the importance of compatibility research in agriculture to assist growers in decisions to co-plant different varieties.

I show that blueberry varieties Emerald, Eureka, Snowchaser, Suziblue, and Twilight can all produce fruit after intra-varietal pollinations. This means that varieties can be planted in solid block plantings and still produce fruit. However, despite Emerald, Eureka and Snowchaser not yielding significantly larger fruit for inter-varietal compared to intra-varietal pollinations, the direction of these differences was always consistent (inter-varietal pollinations always increased adjusted fruit mass) (Fig 1.2). This suggests that the effect sizes may have been too small to pick up using the level of replication in this study and it may take extremely large sample sizes to detect significant treatment differences for some of these varieties. Nevertheless, the consistency of the effect direction (inter-varietal crosses yield greater fruit mass than intra-varietal crosses, Fig. 1.2) suggests that it is prudent to interplant whenever possible. Other studies have also shown that fruit mass can be increased when varieties are pollinated with pollen from other varieties (Bell et al., 2012; Harrison et al., 1993; Müller et al., 2013; Taber and Olmstead, 2016). In this study, the degree of benefit from inter-varietal pollen appeared to differ between varieties and while absolute positive effects were always detected, these were often not statistically significant. For Suziblue and Twilight, inter-varietal pollination also led to significant reductions in developmental time. Inter-varietal pollination has also been shown to decrease developmental time in other blueberry varieties such as Bluecrop, Bluegold, Legacy, Sierra, Toro and Patriot (Dogterom et al., 2000; Ehlenfeldt, 2001; Gupton and Spiers, 1994; Mackenzie, 1997; Müller et al., 2013; Taber and Olmstead, 2016). Choosing the correct pollen source appears to be especially important for varieties such as Eureka, Suziblue and Twilight as pollen quality is likely to have large consequences for yields and interplanting varieties to promote outcrossing therefore seems to be the best strategy for increased fruit mass and decreased developmental times. In contrast, varieties Snowchaser and Emerald appear to be slightly less affected by pollen source and may be better suited to conditions where pollinators seldom move between varieties.

While this study used hand-pollinations to optimise pollination conditions actual pollinators are more likely to deposit mixed pollen loads. Consequently, even with good pollinator movement between varieties, animal pollination is unlikely to achieve the fruit quality recorded in this study. Nevertheless, these results demonstrate that pollinator movement between varieties is likely to increase yields for many commercial blueberry varieties. However, there is little to no data available on how frequently pollinators move between varieties, and how position, spacing and the identity of varieties and pollinator species are likely to affect probabilities of pollen movement between varieties. This represents a fertile area for future study.

While many farms attempt to maximize the chances of pollen movement between cross-compatible varieties by planting different varieties in rows, it is unclear how frequently pollinators actually move between rows. By studying pollinator behaviour, I may improve spacing or placement of blueberry plants, to increase outcrossing. This may be done by matching planting structure with pollinator movement. For example, if a pollinator often moves between rows in a blueberry orchard, then varieties can be planted next to each other in rows because this may improve the chances of pollinator movement from donors to recipients and *vice versa*. However, if pollinators mostly move down rows and varieties are planted next to each other, varieties will be pollinated with pollen from different plants of the same variety. Improving spacing, or placement of blueberry plants may increase the yields currently being achieved under all pollinator environments for any variety requiring inter-varietal pollination to produce optimal yields.

## Acknowledgements

We thank Nicole Vorster, Sarah Vorster, Katya Kapdi and Wesley Hartmann for their help with field collections. Funding was received from National Research Foundation (South Africa), South African Berry Producers Association, Claude Leon Foundation and The Eva Crane Trust.

