## Supplementary material for "Cross-compatibility of five highbush blueberry varieties and ideal crossing combinations": Supplemetary Material

**Table S1:** Comparison of fruit quality variables between different donor pairs for Suziblue as a recipient.

| | SuzC→ SuzC vs TwiC→SuzC | SuzC→ SuzC vs EmC→ SuzC | SuzC→ SuzC vs EurC→ SuzC | EmC→ SuzC vs TwiC→ SuzC | EurC→ SuzC vs TwiC→ SuzC | EurC→ SuzC vs EmC→ SuzC |
|---|---|---|---|---|---|---|
| **Weight** | *z*= 1.991, *p*=0.1862 | ***z*= 3.872, *p*<0.001** | *z*= 0.5940, *p*= 0.9322 | *z*= 2.4050, *p*=0.0736 | *z*= -0.9970, *p*=0.7462 | ***z*= -2.9220, *p*=0.0174** |
| **Developmental period** | *z*= 0.3880, *p*=0.9801 | ***z*= -3.8970, *p*<0.001** | *z*= -0.2760, *p*=0.9926 | ***z*= -4.0920, *p*=<0.001** | *z*= -0.6040, *p*=0.9306 | ***z*= 3.1350, *p*=0.0091** |
| **Fruit set** | *Chi-square*= 6.9299, *df*=3, *p*=0.0742 | | | | | |

Significance indicated by bold type. Arrows indicate the direction of pollen transfer.

**Table S2:** Comparison of fruit quality variables between different donor pairs for Twilight as a recipient.

| | TwiC→TwiC vs SuzC→TwiC | TwiC→TwiC vs EmC→TwiC | TwiC→TwiC vs EurC→TwiC | EmC→ TwiC vs SuzC→TwiC | EurC→TwiC vs SuzC→TwiC | EurC→TwiC vs EmC→TwiC |
|---|---|---|---|---|---|---|
| **Weight** | ***z*= 2.8770, *p*=0.0202** | ***z*= 2.7940, *p*=0.0259** | *z*= 1.5030, *p*=0.4299 | *z*= -0.7010, *p*=0.8946 | *z*= -1.4580, *p*= 0.4573 | *z*= -1.7480, *p*=0.2935 |
| **Developmental period** | ***z*= -5.5930, *p*<0.001** | ***z*= -6.0650, *p*<0.001** | *z*= -0.6950, *p*=0.8990 | *z*= -0.3250, *p*=0.988 | ***z*= 5.1660, *p*<0.001** | ***z*= 5.6580, *p*<0.001** |
| **Fruit set** | *Chi-square*= 3.2940, *df*=3, *p*=0.3485 | | | | | |

Significance indicated by bold type. Arrows indicate the direction of pollen transfer.

**Table S3:** Comparison of fruit quality variables between different donor pairs for Emerald as a recipient.

| | **EmC→ EmC vs. EmC→EurC** |
|---|---|
| **Weight** | *z*=0.7761, *p*=0.4380 |
| **Developmental period** | *z*= 0.9230, *p*=0.3867 |
| **Fruit set** | *Chi-square*=0.1100, *df*=1, *p*=0.7394 |

Significance indicated by bold type. Arrows indicate the direction of pollen transfer.

**Table S4:** Comparison of fruit quality variables between different donor pairs for Eureka as a recipient.

| | **EurC→EurC vs EmC→ EurC** | **EurC→EurC vs SnwC→ EurC** | **EmC→ EurC vs SnwC→ EurC** |
|---|---|---|---|
| **Weight** | *z*=2.0960, *p*=0.0894 | *z*= 1.8350, *p*= 0.1563 | *z*= -0.7130, *p*=0.7539 |
| **Developmental period** | *z*= -0.3120, *p*=0.9480 | *z*= -0.6190, *p*= 0.8090 | *z*= -0.3250, *p*=0.9430 |
| **Fruit set** | *Chi-square*=5.7893, *df*=2, *p*=0.0553 | | |

Significance indicated by bold type. Arrows indicate the direction of pollen transfer.

**Table S5:** Comparison of fruit quality variables between different donor pairs for Snowchaser as a recipient.

| | **SnwC→ SnwC vs EurC →SnwC** |
|---|---|
| **Weight** | *t*=0.9760, *p*=0.3290 |
| **Developmental period** | *t*=-1.0860, *p*= 0.2780 |
| **Fruit set** | *Chi-square*=0.2482, *df*=1, *p*=0.6183 |

Significance indicated by bold type. Arrows indicate the direction of pollen transfer.
